# Evaluation of ChIC-based data requires normalization that properly retains signal-to-noise ratios

**DOI:** 10.1101/2021.08.14.456176

**Authors:** Bofeng Liu, Fengling Chen, Wei Xie

## Abstract

Several chromatin immunocleavage-based (ChIC) methods using Tn5 transposase have been developed to profile histone modifications and transcription factors bindings^1-5^. A recent preprint by Wang et al. raised potential concerns that these methods are prone to open chromatin bias^6^. While the authors are appreciated for alerting the community for this issue, it has been previously described and discussed by Henikoff and colleagues in the original CUT&Tag paper^3,7^. However, as described for CUT&Tag^3^, the signal-to-noise ratio is essential for Tn5-based profiling methods and all antibody-based enrichment assays. Based on this notion, we would like to point out a major analysis issue in Wang et al. that caused a complete loss or dramatic reduction of enrichment at true targets for datasets generated by Tn5-based methods, which in turn artificially enhanced the relative enrichment of potential open chromatin bias. Such analysis issue is caused by distinct background normalizations used towards ChIP-based (chromatin immunoprecipitation) data and Tn5-based data in Wang et al. Only the normalization for Tn5-based data, but not ChIP-seq based data, yielded such effects. Distortion of such signal-to-noise ratio would consequently lead to misleading results.

## Results

Recently, several ChIC methods using Tn5 transposase have been developed to profile histone modifications and transcription factors bindings, including ChIL-seq, CUT&Tag, ACT-seq, CoBATCH, and Stacc-seq^1-5^. Those methods have been applied for studying epigenetic programming and transcription regulation in low-input samples or even single cells^4,5,8,9^. A recent preprint by Wang et al. raised potential concerns that these methods are prone to open chromatin bias^6^. While the authors are appreciated for alerting the community for this issue, it has been previously described and discussed by Henikoff and colleagues in the original CUT&Tag paper^3,7^. For example, such open chromatin bias can be detected in CUT&Tag profiling of NPAT, a transcriptional coactivator of the replication-dependent histone genes that binds only ∼80 genomic sites in the histone gene clusters^3^. Peak calling can identify ∼ 8,600 sites that also include 10% of ATAC-seq defined accessible sites. However, the enrichment of NPAT at histone gene clusters is much stronger than that at non-histone genes and ATAC-seq peaks, and occupies the vast majority of total reads^3^.

We also agree that open chromatin bias remains a potential concern for Tn5-based profiling methods. In our hands, whether open chromatin bias exists and, if so, its extent vary for different antibodies and cell types even using the same protocol. Therefore, extensive tuning of the protocols should be done so that they are tailored for each antibody. However, we would like to emphasize that, as described for CUT&Tag^3^, the target-noise ratio is essential for Tn5-based profiling methods (in fact for all antibody-based experiments). Based on this, it is important to point out a major analysis issue in Wang et al. that caused a complete loss or dramatic reduction of H3K27me3 enrichment at Polycomb targets (“target”) for datasets generated by Tn5-based methods, which in turn artificially enhanced the relative enrichment of potential open chromatin bias (“noise”). Distortion of such signal-to-noise ratio would consequently lead to misleading results. Such analysis issue is caused by the fact that distinct normalizations were used towards ChIP-based (chromatin immunoprecipitation) data and Tn5-based data in Wang et al. Only the normalization for Tn5-based data, but not ChIP-seq based data, yielded such effects. This is shown by detailed re-analysis below.

In Wang et al., benchmarking analysis was performed on H3K27me3 profiles in mESCs generated by conventional ChIP-seq^10^, CoBATCH^5^, CUT&Tag^4^, Stacc-seq^4^, itChIP-seq^11^, and ULI-NChIP^12^ data. The authors used “fold-changes” (FC) between raw reads from “ChIP or ChIC” and a “background” control to evaluate the H3K27me3 enrichment. For ChIP-based methods (conventional ChIP-seq, ULI-NChIP, and itChIP-seq), the fold-changes were calculated over the background **estimated from the corresponding input samples (from fragmented chromatin without going through immunoprecipitation)**. For Tn5-based methods (CUT&Tag, Stacc-seq, and CoBATCH, hereafter referred to as “ChIC”) which do not have input samples, the fold-changes were calculated over the background **estimated from the ChIC sample themselves** (equivalent to immunoprecipitation samples for ChIP-based methods) as described in Methods of Wang et al. (Fig. 1A). Based on the MACS manual^13^, such analysis was done by estimating signals from the IP data by taking the average signals in surrounding 5kb or 10kb regions. It is worth noting that whether the assumption that the nearby 5kb or 10kb regions are true background signals holds true for histone marks such as H3K27me3, which often exist as broad domains, remains to be tested. For convenience, we termed these two FC values as FC-normalized by input (FC-I) and FC-normalized by ChIP/ChIC estimated background (FC-C). FC-C was used in most figures (Fig. 1a-d, Fig. 2, Fig. 3 and Supplementary Fig. 3) in Wang et al. In a few cases (Fig. 1c-d and Supplementary Fig. 2a-b), the fold-change was calculated using background estimated from IgG data for Stacc-seq and CoBATCH (FC-IgG).

**Fig. 1.**
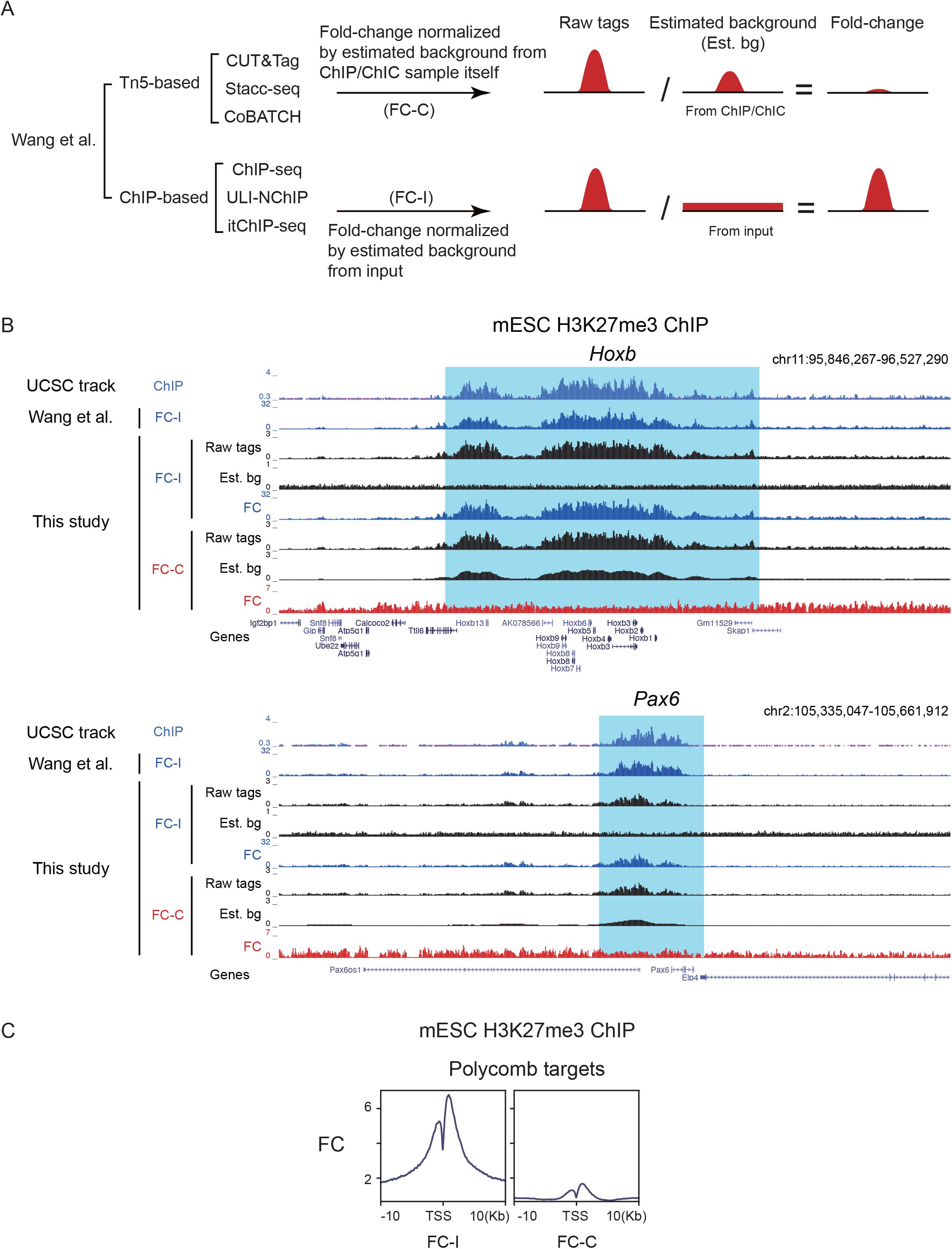
Different normalization between Tn5-based profiling and ChIP-seq in Wang et al. **A**, Schematic illustration of different normalization methods used in Wang et al. **B**, The UCSC browser views showing mESC H3K27me3 ChIP-seq (ENCODE) signal enrichment at *Hoxb* clusters and *Pax6*. **C**, Line charts showing mESC H3K27me3 ChIP-seq enrichment (fold-change normalized by background estimated from input samples (FC-I) or ChIP/ChIC sample itself (FC-C)) at Polycomb targets (Methods).

**Fig. 2.**
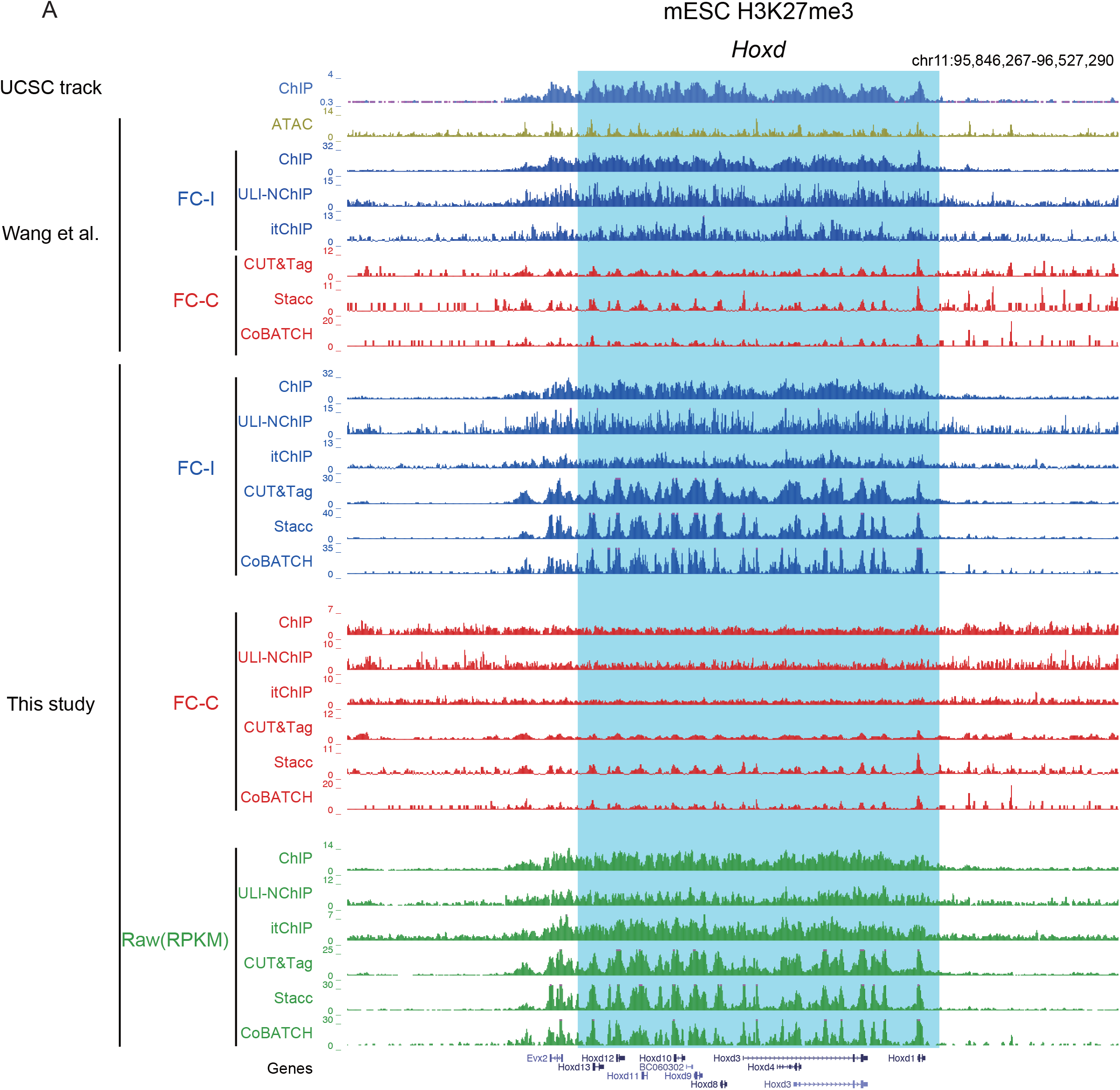
H3K27me3 ChIP-seq or ChIC-based profiling results by different normalization methods at the *Hoxd* cluster. **A**, The UCSC browser views showing mESC H3K27me3 signal enrichment from different normalization and experimental methods at the *Hoxd* cluster. FC-I, fold-change normalized by input samples; FC-C, fold-change normalized by ChIP/ChIC sample itself; RPKM, reads per kilobase per million mapped reads.

**Fig. 3.**
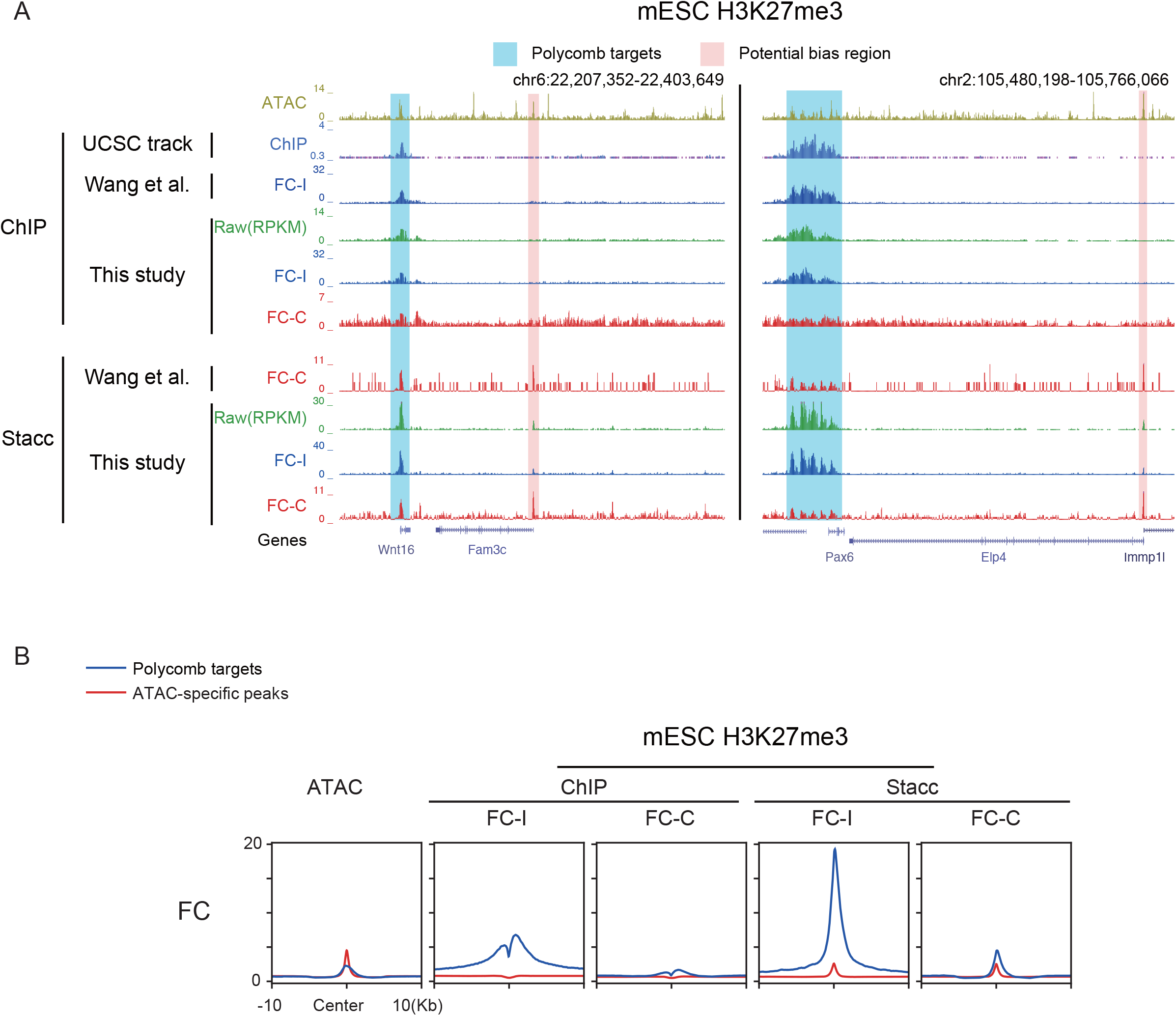
Fold-change normalized by background estimated from ChIP/ChIC sample itself dramatically distorted the signal-to-noise ratios between Polycomb targets and open chromatin. **A**, The UCSC browser views showing mESC ATAC-seq, H3K27me3 ChIP-seq (ENCODE), and Stacc-seq signal enrichment at *Wnt16, Pax6*, and their nearby regions. FC-I, fold-change normalized by input samples; FC-C, fold-change normalized by ChIP/ChIC sample itself; RPKM, reads per kilobase per million mapped reads. **B**, Line charts showing mESC ATAC-seq, H3K27me3 ChIP-seq enrichment (fold-change normalized by background estimated from input samples (FC-I) or ChIP/ChIC sample itself (FC-C)) at Polycomb targets and ATAC-specific peaks (defined by ATAC-seq peaks which are nonoverlapped with ChIP-seq peaks).

We first noticed the normalization difference as the UCSC genome browser snapshot of H3K27me3 Stacc-seq data in Fig. 1b of Wang et al. is dramatically different from the raw tags (RPKM) processed in our lab (Fig. S1, compare Stacc-seq “FC-C” vs. “Raw”). For an unbiased comparison, we performed FC-I and FC-C normalization for all datasets including both ChIP-based and Tn5-based datasets. As Tn5-based datasets do not have “input”, we used IgG data in this analysis. By reproducing the analysis pipeline from Wang et al., we found that even for the same H3K27me3 ChIP-seq reference dataset from ENCODE, FC-I and FC-C values are dramatically different. FC-C, but not FC-I, leads to drastic loss of enrichment at Polycomb targets including the *Hox* regions (Fig. 1B, red, Fig. 1C, right). An expanded analysis showed that the loss of Polycomb target enrichment by FC-C normalization was observed for all ChIP-seq and Tn5-based datasets (Fig. 2, and Fig. S2). This is consistent with the dramatic reduction of H3K27me3 for all FC-C normalized Tn5-based datasets at Polycomb targets in the snapshot in Wang et al. (Fig. 1b in Wang et al.; see Fig. S1). By contrast, strong H3K27me3 enrichment is preserved at Polycomb targets including the *Hox* regions for all FC-I normalized datasets and the raw tags (RPKM), including both those generated by ChIP-based methods and Tn5-based methods (Fig. 1B-C, 2, Fig. S1, S2). We believe that the loss of H3K27me3 in Polycomb targets by FC-C normalization is that it incorrectly takes local broad domains of H3K27me3 signals as background (see Fig. 1B, “Est. bg” track), which automatically cancels out the enrichment at true targets.

Consequently, the loss of H3K27me3 enrichment at Polycomb targets is accompanied by the increase of relative enrichment in potential open chromatin bias regions. For example, the potential open chromatin biased regions of Stacc-seq could be seen near Polycomb target gene *Wnt16* and *Pax6* (Fig. 3A, blue and red shades). The H3K27me3 enrichment of Stacc-seq in Polycomb target gene *Wnt16* and *Pax6* is much stronger than that at nearby potential open chromatin regions with RPKM or FC-I normalized datasets (Fig. 3A, blue vs. red shades, “Raw” and “FC-I”). However, such differences are clearly erased in FC-C normalized datasets (Fig. 3A, blue vs. red shades, “FC-C”). This observation is further confirmed by a global average analysis, as the enrichment of ChIP and Stacc-seq, based on FC-I, at Polycomb targets is markedly higher than that in open chromatin regions without H3K27me3 (Fig. 3B, “FC-I”). However, such difference was largely eliminated after FC-C normalization (Fig. 3B, “FC-C”). In sum, we believe that the benchmarking analyses in Wang et al. implemented differential analyses for Tn5-based data (FC-C) and ChIP based data (FC-I), and the FC-C normalization for Tn5-based data particularly causes a dramatic reduction of H3K27me3 enrichment at Polycomb targets and the “signal-to-noise” ratios, therefore artificially increasing the relative enrichment of open chromatin bias.

Finally, we note that in our hands, whether an antibody can effectively capture epitopes, whether it produces open chromatin bias (for Tn5-based approaches), and (if exists) its extent vary widely, which likely depends on the affinity of different antibodies (even for the same epitope), the abundance of epitopes, and the local chromatin environment of target proteins. We disagree that a fixed experimental condition and protocol should be used for all experiments. Given the diverse nature of different antibodies, cell types and experimental conditions, we suggest that the specific experimental conditions such as the wash step should be tuned for different antibodies in a case-by-case manner, as described in our paper^4^. Choices of methods (including other antibody-based approach such as CUT&RUN^14^) should be determined based on the actual performance toward each antibody-epitope pair, and appropriate experimental controls (such as using cells with target genes mutated or proteins degraded) should be included whenever they are feasible. In our previous study^4^, we validated the binding data of Pol II subunit Rpb1 using extended inhibitor treatment induced Pol II degradation.

## Discussion

Several ChIC-based methods have been developed in recent years and become increasingly popular in transcription regulation research^1-5^. Despite their high sensitivity and convenient procedure, caution should be taken to rigorously evaluate their pros and cons including the potential open chromatin bias, as illustrated in the original CUT&Tag^3^ paper. However, the extent of such potential bias should be properly evaluated. Based on our analyses, a normalization analysis in Wang et al. resulted in strongly distorted signal-to-noise ratio, a metric that is essential to evaluate the performance of all antibody-based enrichment assays and to correctly interpret the results. Similar to experimental methods, different computational methods should be applied with caution and proper validation. A normalization method used for TF or one type of epigenetic mark needs to be validated before being applied to another epigenetic mark. Finally, we suggest that choices of methods should be experimentally determined based on the actual performance toward each specific antibody-epitope pair with appropriate controls.

## Author Contributions

B.L. and W.X. conceived and designed the project. B.L. and F.C. analyzed data. B.L. and W.X. wrote the manuscript with help from F.C.

## Competing interests

The authors declare no competing interests.

## Acknowledgments

We thank members of the Xie laboratory for discussions and comments on the manuscript. We thank M.W. and Y.Z. for sharing the scripts and data tracks. This work was supported by the National Natural Science Foundation of China (31988101 to W.X.) and the THU-PKU Center for Life Sciences (W.X.). B.L. is supported by postdoctoral fellowships from the Tsinghua Shuimu Scholar. F.C. is supported by postdoctoral fellowships from the Tsinghua-Peking Joint Center for Life Sciences. W.X. is an HHMI international research scholar.

## Figure legends

**Fig. S1.**
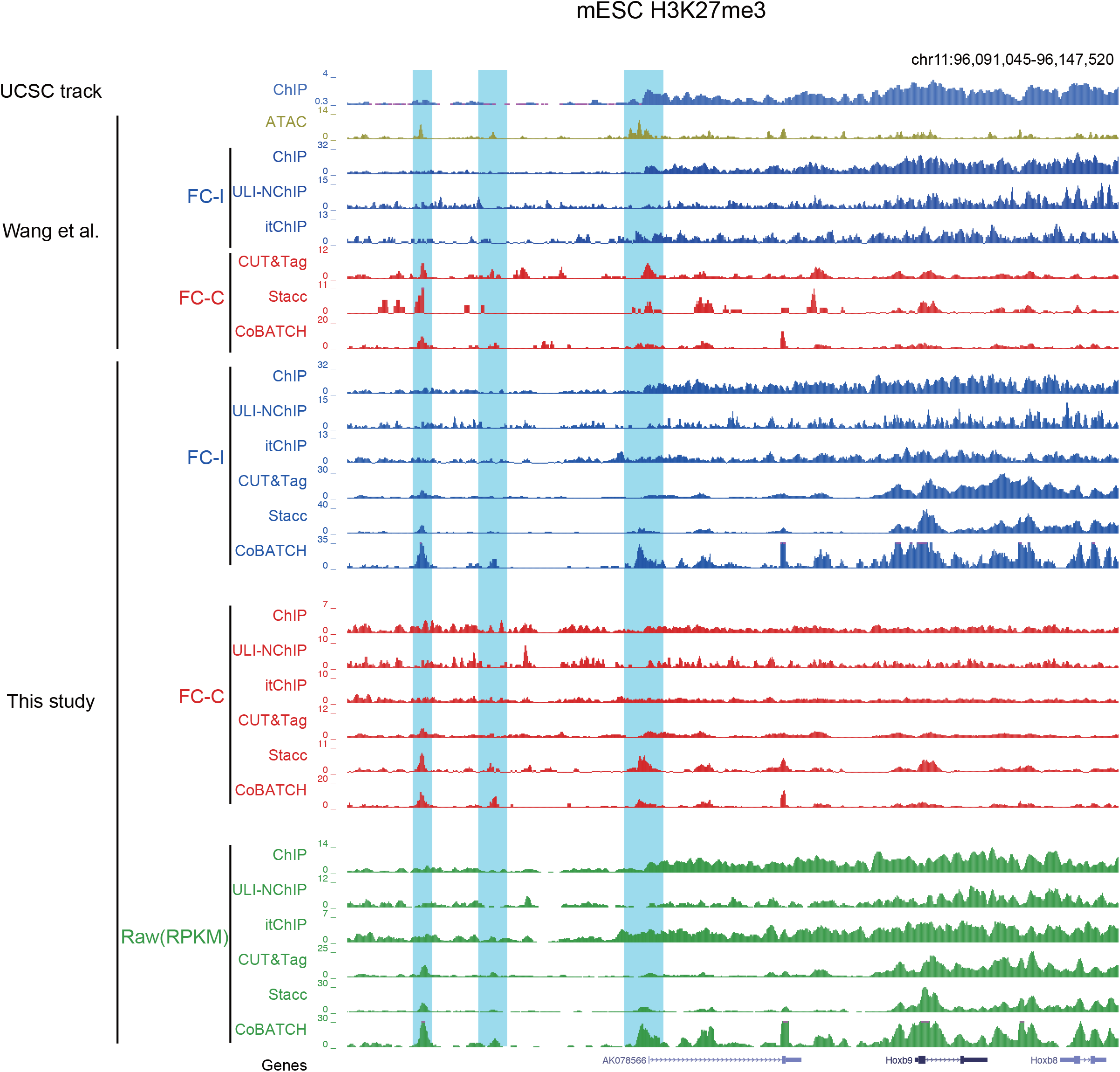
H3K27me3 ChIP-seq or ChIC-based profiling results by different normalization methods in the same region presented in Fig. 1b, Wang et al. **A**, The UCSC browser views showing mESC H3K27me3 signal enrichment from different normalization and experimental methods in the region presented in Fig. 1b, Wang et al. FC-I, fold-change normalized by input samples; FC-C, fold-change normalized by ChIP/ChIC sample itself; RPKM, reads per kilobase per million mapped reads.

**Fig. S2.**
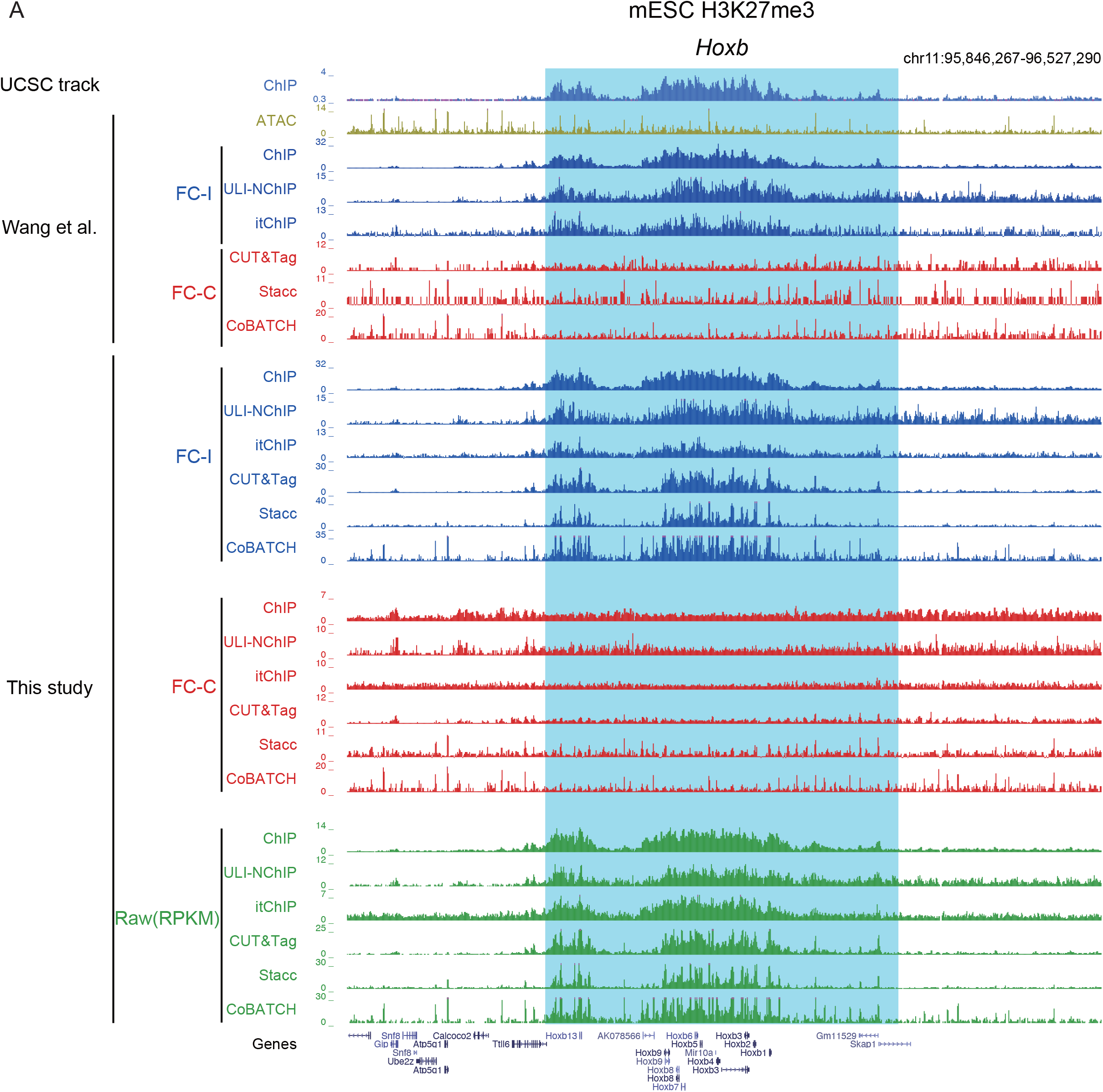
H3K27me3 ChIP-seq or ChIC-based profiling results by different normalization methods at the *Hoxb* cluster. **A**, The UCSC browser views showing mESC H3K27me3 signal enrichment from different normalization and experimental methods at the *Hoxb* cluster. FC-I, fold-change normalized by input samples; FC-C, fold-change normalized by ChIP/ChIC sample itself; RPKM, reads per kilobase per million mapped reads.

## Methods

### Data collection

The same mESC H3K27me3 and ATAC-seq data as Wang et al. (also summarized in Supplementary Table 1) were used. The data tracks marked by Wang et al. were shared by Wang et al. and were transformed from mm10 to mm9 by LiftOver.

### ChIP, ChIC, and ATAC-seq data processing

The single-end reads were aligned to mm9 with the parameters: -t -q -N 1 -L 25 by Bowtie (version 2.2.2)^15^. The paired-end reads were aligned to mm9 with the parameters: -t -q -N 1 -L 25 -X 2000 --no-mixed --no-discordant by Bowtie (version 2.2.2)^15^. All unmapped reads, non-uniquely mapped reads, and PCR duplicates were removed.

### Peak calling and track generation

All the peaks were called by MACS2 with the parameters -B –SPMR -p 1e-4 -g mm --broad -- broad-cutoff 1e-4 --keep-dup all --scale-to large, following the method described in Wang et al. The parameter -c was applied when the input control sample was included, which refers to FC-normalized by input (FC-I). Peak calling without -c was used when the input control sample was not included, which refers to FC-normalized by ChIP/ChIC estimated background (FC-C).

The fold-change signal tracks were generated using MACS2 bdgcmp with the input of treat-pileup and control-lambda bedgraph files generated from MACS2 callpeak in the last step.

The RPKM signal tracks were generated by computing the numbers of reads per kilobase of bin per million of reads sequenced (RPKM). To visualize the signal in the UCSC genome browser, we extended each read by 250 bp and counted the coverage for each base.

### Peak comparison, TSS analysis, and Polycomb target analysis

Peaks were compared by BEDTools (version 2.27.1)^16^. Peaks with at least half-length intersected were considered as overlapping (parameter -f 0.5 in intersectBed). The TSSs were from annotated promoters (RefSeq, Ensemble, and UCSC Known Gene databases combined). Polycomb targets were from Liu et al.^4^. Briefly, genes marked by H3K27me3 at their promoters (+/- 5kb) in mESCs (based on ChIP-seq) were termed as Polycomb targets.

## Code availability

Software and code used to analyze these data are available at GitHub: https://github.com/BofengLiu/Xielab

## Data availability

The public datasets are summarized in Supplementary Table 1.

## Notes

https://github.com/BofengLiu/Xielab

### Competing Interest Statement

The authors have declared no competing interest.

